# Uncovering the genetic architecture and evolutionary roots of androgenetic alopecia in African men

**DOI:** 10.1101/2024.01.12.575396

**Authors:** Rohini Janivara, Ujani Hazra, Aaron Pfennig, Maxine Harlemon, Michelle S. Kim, Muthukrishnan Eaaswarkhanth, Wenlong C. Chen, Adebola Ogunbiyi, Paidamoyo Kachambwa, Lindsay N. Petersen, Mohamed Jalloh, James E. Mensah, Andrew A. Adjei, Ben Adusei, Maureen Joffe, Serigne M. Gueye, Oseremen I. Aisuodionoe-Shadrach, Pedro W. Fernandez, Thomas E. Rohan, Caroline Andrews, Timothy R. Rebbeck, Akindele O. Adebiyi, Ilir Agalliu, Joseph Lachance

**Affiliations:** School of Biological Sciences, Georgia Institute of Technology, Atlanta, Georgia, USA; Department of Biology, Morgan State University, Baltimore, Maryland, USA; Department of Human Genetics University of Michigan, Ann Arbor, Michigan, USA; Strengthening Oncology Services Research Unit, Faculty of Health Sciences, University of the Witwatersrand, Johannesburg, South Africa; Sydney Brenner Institute for Molecular Bioscience, Faculty of Health Sciences, University of the Witwatersrand, Johannesburg, South Africa; National Cancer Registry, National Institute for Communicable Diseases a Division of the National Health Laboratory Service, Johannesburg, South Africa; College of Medicine, University of Ibadan, Ibadan, Nigeria; Centre for Proteomic and Genomic Research, Cape Town, South Africa; Mediclinic Precise Southern Africa, Cape Town, South Africa; Université Cheikh Anta Diop de Dakar, Dakar, Senegal; Université Iba Der Thiam de Thiès, Thiès, Senegal; Korle-Bu Teaching Hospital and University of Ghana Medical School, Accra, Ghana; Department of Pathology, University of Ghana Medical School, Accra, Ghana; 37 Military Hospital, Accra, Ghana; College of Health Sciences, University of Abuja, University of Abuja Teaching Hospital and Cancer Science Centre, Abuja, Nigeria; Faculty of Medicine and Health Sciences, Stellenbosch University, Cape Town, South Africa; Department of Epidemiology and Population Health, Albert Einstein College of Medicine, Bronx, New York, USA; Dana-Farber Cancer Institute, Boston, Massachusetts, USA; Harvard T.H. Chan School of Public Health, Boston, Massachusetts, USA

**Keywords:** Africa, evolutionary genetics, GWAS, male pattern baldness, polygenic risk scores

## Abstract

Androgenetic alopecia is a highly heritable trait. However, much of our understanding about the genetics of male pattern baldness comes from individuals of European descent. Here, we examined a novel dataset comprising 2,136 men from Ghana, Nigeria, Senegal, and South Africa that were genotyped using a custom array. We first tested how genetic predictions of baldness generalize from Europe to Africa, finding that polygenic scores from European GWAS yielded AUC statistics that ranged from 0.513 to 0.546, indicating that genetic predictions of baldness in African populations performed notably worse than in European populations. Subsequently, we conducted the first African GWAS of androgenetic alopecia, focusing on self-reported baldness patterns at age 45. After correcting for present age, population structure, and study site, we identified 266 moderately significant associations, 51 of which were independent (p-value < 10^-5^, r^2^ < 0.2). Most baldness associations were autosomal, and the X chromosomes does not appear to have a large impact on baldness in African men. Finally, we examined the evolutionary causes of continental differences in genetic architecture. Although Neanderthal alleles have previously been associated with skin and hair phenotypes, we did not find evidence that European-ascertained baldness hits were enriched for signatures of ancient introgression. Most loci that are associated with androgenetic alopecia are evolving neutrally. However, multiple baldness-associated SNPs near the *EDA2R* and *AR* genes have large allele frequency differences between continents. Collectively, our findings illustrate how evolutionary history contributes to the limited portability of genetic predictions across ancestries.

## Introduction

Most complex traits are polygenic, and their genetic architectures have been shaped by evolutionary history.^1,2^ Phenomena such as population bottlenecks and geographical isolation can cause the relative importance of different genetic risk factors to vary by population.^3,4^ Similarly, natural selection can also impact population-specific genetic architectures.^5–7^ Because of this, genetic predictors that are developed in one part of the world often transfer poorly to other parts of the world.^8–10^ However, the portability of genetic predictions can be trait-specific.^11^

Androgenetic alopecia, commonly known as male-pattern baldness (MPB), is a model trait due to its variable prevalence across populations, relevance to health, and high heritability. This condition typically involves progressive loss of hair above the temples and at the vertex of the scalp due to the progressive miniaturization of hair follicles.^12^ Notably, hair morphology varies by ancestry,^13^ and the prevalence of androgenetic alopecia varies across the world. The onset of MPB tends to be later in Japanese men, and men of non-European descent are more likely to retain their frontal hair lines than men of European descent.^14^ By contrast, individuals of Asian descent have higher rates of alopecia areata, an inflammatory condition of hair follicles that results in patchy bald spots.^15^ Disparities also exist for other forms of alopecia, including scarring alopecia (i.e., CCCA), which is disproportionally observed in women of African descent.^16^ Importantly, baldness is not only a cosmetic trait – it has also been associated with endocrine, metabolic, and cardiovascular diseases, likely due to the myriad effects of sex hormones.^17^ Baldness has long been associated with heredity,^18^ and men whose fathers and/or maternal grandfathers are bald are more likely to experience hair loss.^19^ Heritability estimates of MPB range from 60% to 70%,^20^ and genetic risk factors in European populations include polymorphisms at 20p11 and Xq12, including alleles near the *EDA2R* (ectodysplasin A2 receptor) and *AR* (androgen receptor) genes.^21,22^ Intriguingly, introgressed DNA from Neanderthals is enriched for genes that are involved in keratin formation, i.e., skin and hair phenotypes.^23,24^ Because Neanderthal DNA is almost exclusively found in non-African genomes,^25^ ancient introgression may contribute to continental differences in the genetic architecture of MPB.

During the past few years several large-scale genome-wide association studies (GWAS) have expanded our understanding of the genetic architecture of MPB. In an analysis of over 22,000 European men, Heilmann-Heimbach et al. conducted the first large meta-analysis of MPB.^26^ European individuals in the top quartile of a polygenic score (PGS) built from this study were more likely to be bald than individuals in the lowest quartile (odds ratio = 4.16, 63 independent GWAS loci).^26^ In a study of over 70,000 men, Piratsu et al. identified 71 susceptibility loci that were enriched for *Wnt* signaling and apoptosis pathways.^27^ In a study of over 52,000 participants from the UK Biobank, Hagenaars et al. identified over 270 independent autosomal and X-linked variants associated with hair loss.^28^ The Hagenaars et al. PGS using these variants was able to distinguish between European men who had severe hair loss vs. no hair loss for prediction models that included age (AUC = 0.78).^28^ A more recent study by Chen et al., that used feature selection prior to training a polygenic predictor on the UK Biobank dataset, reported slightly boosted AUC statistics of severe hair loss vs. no hair loss in European populations (AUC = 0.81 when age was included).^17^ MPB-associated loci from the UK Biobank are enriched for weak signatures of negative selection, perhaps due to pleiotropic effects on other traits.^29^ Additional analyses of over 72,000 exomes from the UK Biobank suggest that rare variants make only a small contribution to the overall risks of MPB in Europe.^30^ However, one limitation of these large-scale genetic studies is that they largely focused on European cohorts, thereby skewing our understanding of variants that contribute to MPB. Consequently, it is unknown if the genetic architecture of MPB differs for African men.

To overcome existing knowledge gaps, we tested how well European-ascertained polygenic risk scores for male-pattern baldness generalize to sub-Saharan African populations. We then inferred the population genetic architecture of this complex trait by performing the first African GWAS of androgenetic alopecia. Our subsequent analyses explored multiple evolutionary hypotheses for the poor portability of genetic predictions. These potential causes include private alleles that are due to recent mutations, Neanderthal alleles that are only informative about baldness in Europeans, and natural selection acting on baldness-associated loci.

## Subjects and Methods

### Baldness phenotyping

African participants included in this study were men without a diagnosis of prostate cancer or any other cancers who were recruited as controls for a large case-control study by the Men of African Descent and Carcinoma of the Prostate (MADCaP) Network^31,32^ at seven study sites: Hôpital Général de Grand Yoff/Institut de Formation et de Recherche en Urologie in Dakar, Senegal; 37 Military Hospital in Accra, Ghana; Korle-Bu Teaching Hospital in Accra, Ghana; University College Hospital in Ibadan, Nigeria; University of Abuja Teaching Hospital in Abuja, Nigeria; Wits Health Consortium/National Health Laboratory Services in Johannesburg, South Africa; and Stellenbosch University in Cape Town, South Africa. At each study site, participants were recruited using protocols approved by that site’s Institutional Review Board/Ethics Review Board and the central data management center at the Dana-Farber Cancer Institute. Participants completed an interviewer-administered questionnaire that queried them about their hair patterns at age 30 and at age 45 using the Hamilton-Norwood baldness scale. To better integrate our data with previous work (e.g., data field 2395 in the of UK Biobank data) and facilitate an ordinal GWAS of androgenetic alopecia, we rescaled Hamilton-Norwood scores into four different categories: no hair loss, slight hair loss, moderate hair loss, severe hair loss (Figure 1). We then excluded all individuals who had incongruent baldness scores (i.e., more hair at age 45 than at age 30). To avoid the potential for reporting bias and/or misclassification, our subsequent analyses focus on self-reported baldness scores at age 45, while using current age to correct for recall errors. Table S1 contains a full list of the number of individuals per study site in each baldness class and Figure S1 is a PCA plot showing the population genetic structure of this dataset.

**Figure 1.**
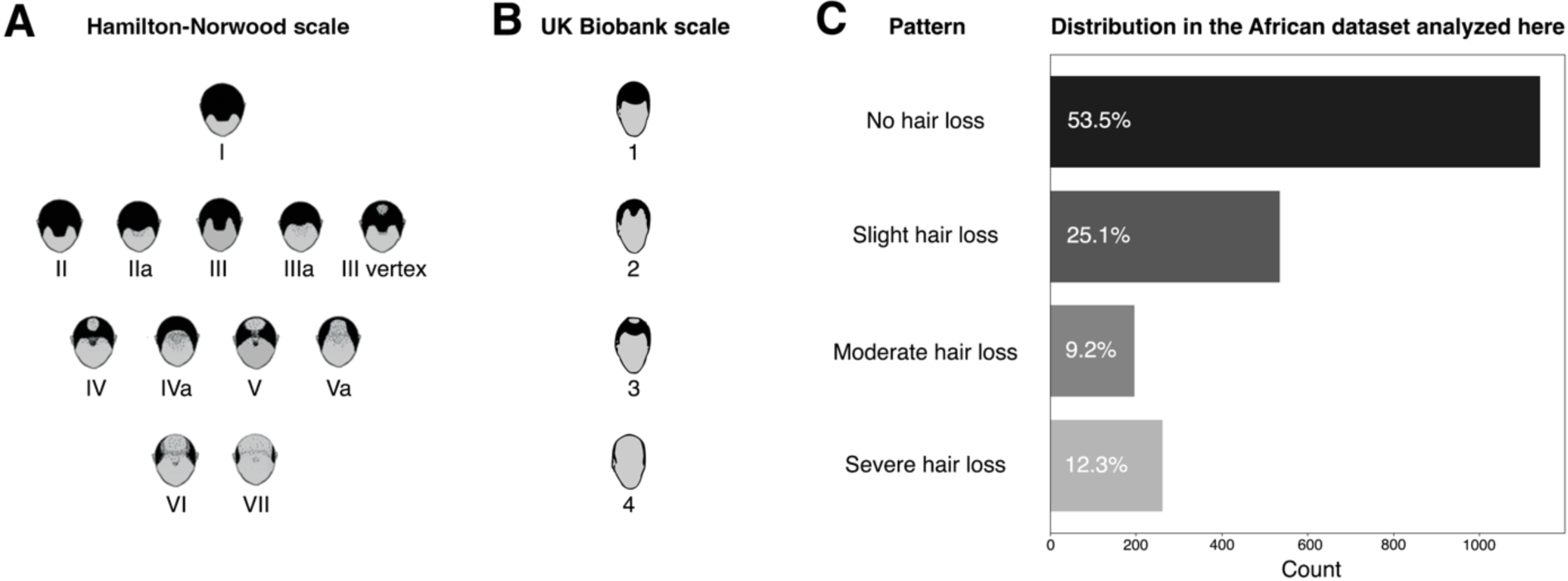
Baldness phenotypes. (A) The Hamilton-Norwood baldness scale partitioned into four categories. (B) Baldness scale from question 2395 of the UK Biobank questionnaire. (C) Proportions of each baldness pattern in the African dataset that was analyzed here. Individuals were sampled from Senegal, Ghana, Nigeria, and South Africa. Study-site specific counts of each baldness phenotype can be found in Table S1.

### Genotyping, QC, and imputation

Genotype data were obtained for African men using MADCaP Array,^33^ a custom genotyping platform that is optimized for detecting genetic associations in sub-Saharan Africa.^33^ Details about SNP calling can be found in other MADCaP Network publications.^33,34^ We performed standard quality control procedures using PLINK. Samples were included if call rates exceeded 98%, they were not related to other samples (kinship coefficient < 0.0884), and they had an African ancestry percentage > 70% (as per previous work^34^). Markers were included if genotype missingness was less than 5%, a minor allele frequency filter was passed (MAF > 0.01), and there were no detectable departures from Hardy-Weinberg proportions (p-value < 10^-5^). After these QC filters, 2,136 unrelated samples with 1,092,609 markers remained. We then imputed additional markers using the TOPMed Imputation Panel and Server.^35^ Post-imputation QC filtering included imputation quality (r^2^ score > 0.8) and minor allele frequency cutoffs (MAF > 0.01), yielding a total of 15,378,257 autosomal and 583,558 X chromosome polymorphisms.

### Tests of polygenic score performance

To assess the portability of genetic predictors of MPB we extracted lists of baldness-associated SNPs from the Hagenaars et. al.^28^ PGS and the Heilmann-Heimbach et. al.^26^ PGS. Using the allele dosage and effect sizes of each trait-associated SNP, we calculated PGS for 2,136 African men. After correcting for covariates the prediction accuracy of each predictor was estimated using the nonparametric Bayesian inference of the covariate-adjusted ROC curve (AROC) package in R.^36^ The Area Under the Curve (AUC) statistics reported here isolate the effects of genetics regarding baldness predictions and they correct for present age, sample site, and population structure (the first three PCs in PCA). Additional tests of PGS performance involved comparing the relative proportions of individuals in each phenotypic class for individuals in the top 10% each PGS distribution compared to individuals in the middle 20% of each PGS distribution. These analyses were repeated for both the Hagenaars et al. PGS and the Heilmann-Heimbach et al. PGS. We note that self-identified white British individuals in the UK Biobank who are aged 40-50 have the following distribution of baldness scores: pattern 1: 31.8%, pattern 2: 23.0%, pattern 3: 26.8%, and pattern 4: 18.4%.

### GWAS analyses

We performed an ordinal model GWAS of MPB, focusing on the four ordered phenotypes in Figure 1: no hair loss, slight hair loss, moderate hair loss, and severe hair loss (i.e., baldness scores range from 1 to 4, as pe question 2395 of the UK Biobank questionnaire). The Proportional Odds Logistic Mixed Model (POLMM) package^37^ was used to estimate p-values and effect sizes of each genetic variant in our ordinal GWAS. The GWAS was corrected for multiple covariates: current age (to correct for errors in self-reported baldness phenotypes), recruitment study site (to correct for possible batch effects in recruitment and referral bias), and the first three principal components (to correct for population structure, Figure S1). LDlink^38^ was used to identify if African GWAS hits were in linkage disequilibrium with previous baldness hits. We used the LDexpress tool^39^ to examine whether any of our African associations with MPB are in LD with GTEx eQTLs, i.e., variants that affect gene expression (thresholds: r^2^ ≥ 0.2, GTEx p-value < 10^-10^, genomic window: ± 500kb). The gene-set enrichment analysis was performed using snpXplorer, a web-server application.^40^ This analysis used the following gene-sets: GO:BP, KEGG, Reactome Wiki Pathways.

The total SNP heritability of MPB for each chromosome was inferred using a modified version of the Linkage Disequilibrium Adjusted Kinships (LDAK)^41^ approach, specifically Basic Protocol 2 as described by Srivastava et al.^42^ This protocol included additional minor allele frequency (MAF ≥ 0,05) and LD-pruning (--indep-pairwise 100 50 0.1) filters in PLINK, as well as pruning out individuals with crypted relatedness using a genetic relatedness matrix (GRM) generated from autosomal data (max relatedness = 0.05). SNP heritability was calculated using REML for chromosomes 1 to 22 together using 22 GRMs, while SNP heritability for the X chromosome was computed separately. This information was then used to determine the relative contribution of each chromosome to the genetic architecture of MPB in Africa by dividing the summed SNP heritability for each chromosome by the total genome-wide SNP heritability.

### Evolutionary genetics analyses

Tests of enrichment of Neanderthal DNA used Skov’s introgression map, which was originally generated from 27,566 Icelandic genomes.^43^ Using this introgression map, we calculated the proportion of Neanderthal DNA in Europeans per non-overlapping 10kb windows across the genome. We then intersected MPB-associated variants in the Hagenaars et. al.^28^ PGS and the Heilmann-Heimbach et. al.^26^ PGS with the tiled map of introgression frequencies. These introgression frequencies were then compared to the distribution of introgression frequencies of 1,000 sets of control SNPs that were matched for allele frequency, local linkage disequilibrium, and distance to the nearest gene. The LiftOver tool was used to harmonize hg19 and hg38 genomic positions.

Population genetic analyses leveraged genetic data from the 1000 Genomes Project (1KGP). Joint site frequency spectrum plots were generated using individuals of African ancestry (ACB, ASW, ESN, GWD, LWK, MSL, and YRI) and European ancestry (CEU, FIN, IBS, GBR, and TSI). A full list of 1KGP population codes can be found at: http://ftp.1000genomes.ebi.ac.uk/vol1/ftp/README_populations.md. Tests of polygenic selection focused on integrative haplotype scores (iHS) of autosomal variants.^44^ A full description of this approach has been described elsewhere.^45^ Briefly, this involved inferring whether sets of MBP-associated SNPs are more likely to have outlier values of iHS statistics (iHS < -1.96 or iHS > 1.96) than 1000 sets of matched control SNPs. F_ST_ calculations used the following equation: *F*_ST_ = *Var*(*p*)⁄(*p̅*(1 − *p̅*)) where *Var*(*p*) refers to the variance in allele frequencies across Europe and Africa and *p̅* refers to the mean allele frequency across both continental populations. These calculations excluded admixed individuals, i.e., African frequencies were generated from ESN, GWD, LWK, MSL, and YRI individuals, while European frequencies were generated from CEU, FIN, IBS, GBR, and TSI individuals. Continental frequencies of haplotypes at Xq12 were calculated from 1KGP data using genotype data at three SNPS: rs2497911, rs12558842, rs1204041.

## Results

### Poor portability of baldness predictions to sub-Saharan Africa

Although multiple polygenic predictors of MBP exist, they have mostly been ascertained in European populations. To find out whether these genetic predictors are informative regarding MPB in Africa, we applied two different PGS to the MADCaP Network dataset. Although the Hagenaars et al. PGS performed well when tested on a European ancestry cohort,^28^ it had only a limited ability to distinguish between any hair loss vs. no hair loss in African men (Figure 2A, AUC = 0.546, 95% CI: 0.522-0.572). Indeed, it performed poorly for slight hair loss vs. no hair loss (AUC = 0.533, 95% CI: 0.503-0.562), moderate hair loss vs. no hair loss (AUC = 0.558, 95% CI: 0.514-0.601), and severe hair loss vs. no hair loss (AUC = 0.565, 95% CI: 0.527-0.603). Despite these low AUC statistics, individuals in the top 10% of the Hagenaars et al. PGS distribution were 33% more likely to have severe baldness at age 45 than individuals in the middle 20% of the PGS distribution (Figure 2B). Similarly, although the Heilmann-Heimbach et al. PGS also performed reasonably well on a European cohort,^26^ its ability to distinguish between any hair loss vs. no hair loss was effectively no better than chance when applied to African men (Fig 2C, AUC = 0.513, 95% CI: 0.488-0.538), and it performed poorly for slight hair loss vs. no hair loss (AUC = 0.504, 95% CI: 0.474-0.533), moderate hair loss vs. no hair loss (AUC = 0.538, 95% CI: 0.496-0.582), and severe hair loss vs. no hair loss (AUC = 0.516, 95% CI: 0.477-0.555). However, individuals in the top 10% of the Heilmann-Heimbach et al. PGS distribution were 50% more likely to be severely bald at age 45 than individuals in middle 20% of the PGS distribution (Figure 2D). Overall, our results reveal that the genetic predictions of androgenetic alopecia developed from European populations do not transfer well to sub-Saharan Africa.

**Figure 2.**
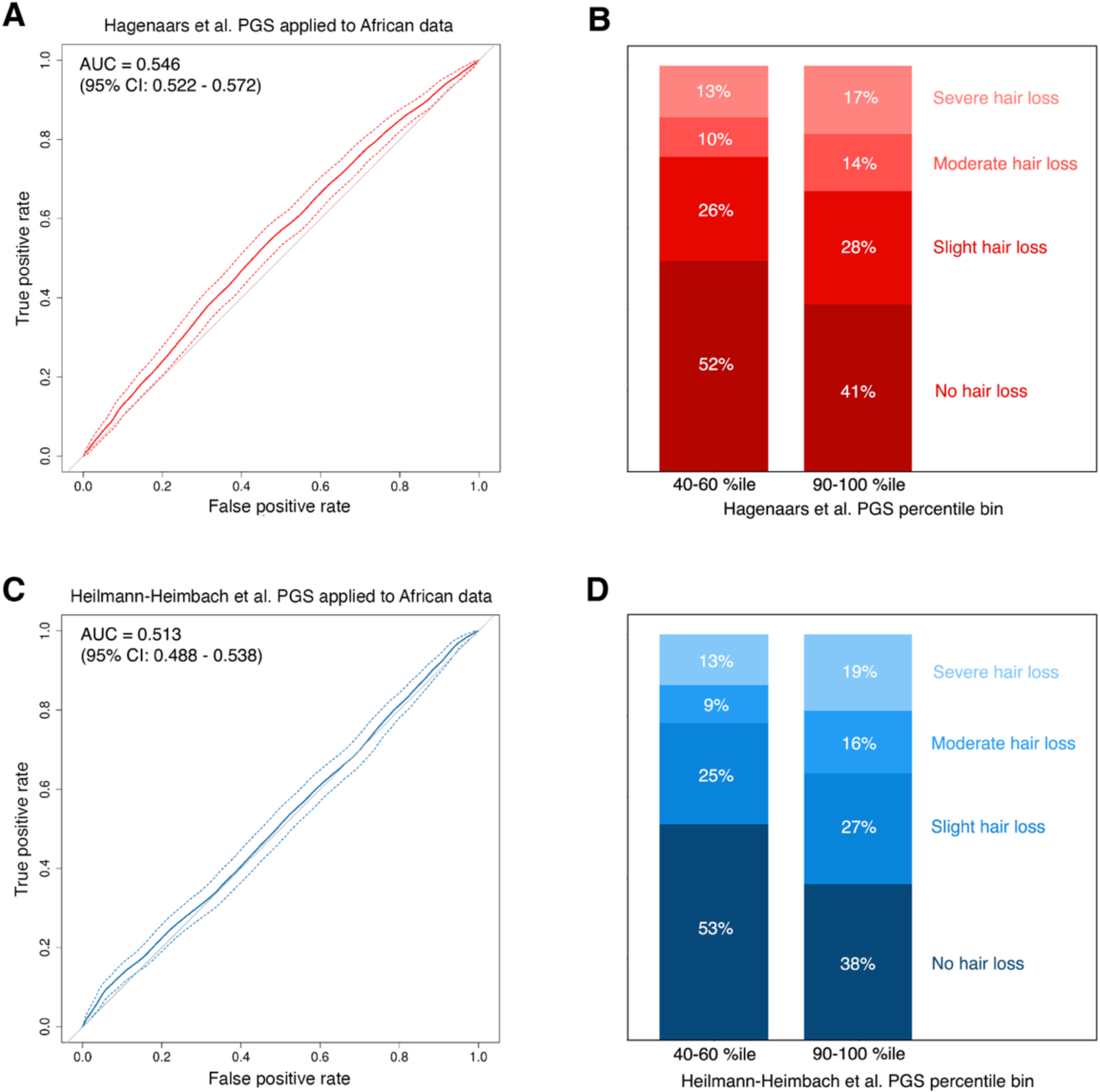
Performance of polygenic scores in Africa. ROC curves focus on comparisons between individuals who have any baldness vs. no baldness. AUC statistics corrected for present age, study site, and population structure. (A) AUC statistics of the Hagenaars et al. PGS applied to African data. (B) Baldness patterns of African men in the middle 20% and top 10% of the Hagenaars et al. PGS distribution. (C) AUC statistics of the Heilmann-Heimbach et al. PGS applied to African data. (D) Baldness patterns of African men in the middle 20% and top 10% of the Heilmann-Heimbach et al. PGS distribution.

### The first African GWAS of male-pattern baldness

Given the poor portability of genetic predictors from European populations, we performed a GWAS of MPB in Africa to identify continental differences in the genetic architecture of this complex trait (Figure 3A). For this we used the MADCaP dataset which contains 2,136 samples from Senegal, Ghana, Nigeria, and South Africa. This GWAS used a proportional odds logistic mixed model which grouped the different intensities of baldness into four ordinal classes. Following adjustments for population structure, study site, and genotype array technology, the genomic inflation factor in this analysis was minimal (λ_GC 0.5_ = 0.998, Figure 3B), indicating the absence of batch effects and confounding. Although our sample size lacked the statistical power to capture any genome-wide significant associations, we found 51 independent marginally significant associations (p-value < 10^-5^; LD pruning threshold: r^2^ < 0.2, Figure 3). A full list of these African hits can be found in Table S2. Among the 51 marginal associations, 36 are not in linkage disequilibrium with previously reported GWAS hits for any trait (r^2^ < 0.2). Key loci that were associated with MPB in Africa include 7q22.2, 1p13.2, and Xq12 (Figure 3A). The 7q22.2 locus contains moderately significant variants in the intronic regions of *COG5* and *HBP1* genes. The lead African SNP at the 1p13.2 locus (rs116494345) is monomorphic in Europe and Asia, i.e., its impact on MPB is Africa-specific. The lead SNP at the Xq12 locus (rs1204041) is in the intronic region of the *AR* (androgen receptor) gene. Other notable baldness associations are rs191219783 (proximal to *CCR7*, a chemokine receptor gene that is expressed by dendritic cells in hair follicles^46^) and rs143451223 (in *SLC301A10*, a manganese transporter gene).

**Figure 3.**
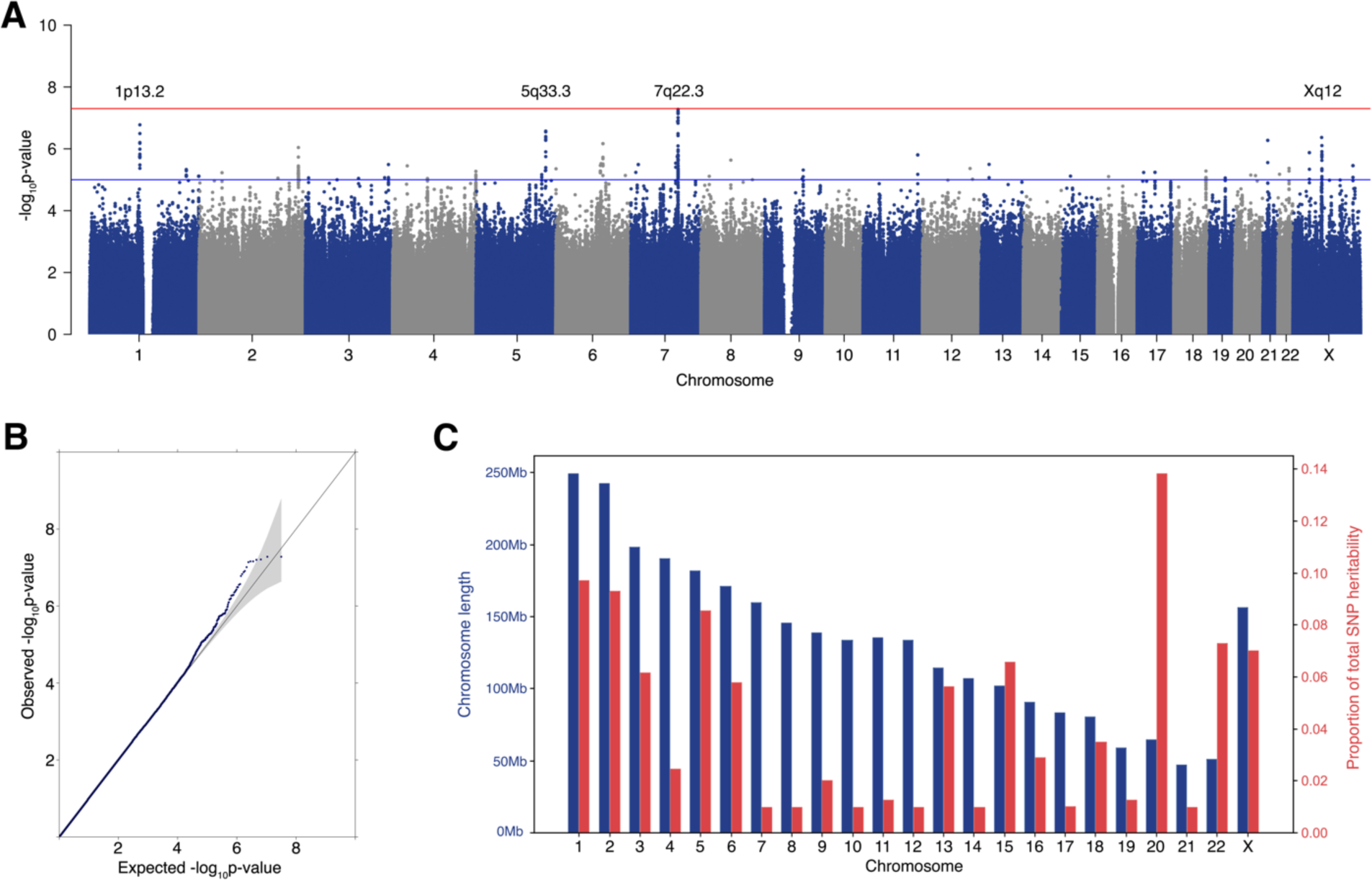
African GWAS of androgenetic alopecia. (A) Manhattan plot for the ordinal GWAS of MPB. Phenotypes were scored using a four-point scale. Sample size: 2,136 African men. (B) QQ plot for the ordinal GWAS of MPB in Africa (λ_GC 0.5_ = 0.998). (C) Physical sizes (blue) and relative contribution to SNP heritability of each chromosome.

We further explored the genetic architecture of MPB in Africa by testing for enrichment of chromosomes, pathways, and tissues. To test whether the X chromosome makes a disproportionally large contribution to the heritability of MPB, we compared relative estimates of the total SNP-heritability to chromosome size. We note that larger chromosomes need not make larger contributions to variation in MPB (Figure 3C). This finding suggests that the genetic architecture of this trait is not extremely polygenic, at least compared to traits like height, where heritability is distributed relatively evenly across the genome.^47^ Interestingly, the X chromosome contribute only modestly to the genetic variance of MBP in African men, i.e. it does not make an outsized contribution to the genetic variance of this complex trait (Figure 3C). Although this pattern likely reflects the underlying biology, we note that lower statistical power to detect X-linked associations compared to autosomal associations may also play a role. Functional annotation of private alleles tends to be limited, especially when these polymorphisms are Africa-specific. One byproduct of this is that 37 of our African associations with MPB are not in linkage disequilibrium with GTEx variants that affect gene expression. Of the remaining 14 African associations, five are in linkage disequilibrium with skin eQTLs and three are in linkage disequilibrium with testis eQTLs. We did not find evidence that the set of 51 African MPB associations were enriched for any specific biological pathway (Table S3).

### Evolutionary genetics of male-pattern baldness

We investigated multiple evolutionary hypotheses to explain the poor portability of polygenic predictions and continental differences in the genetic architecture of MPB. Possible explanations involve the presence of private alleles, ancient introgression of Neanderthal DNA, and natural selection acting on genomic regions that have been associated with baldness. Comparisons between the allele frequencies of baldness-associated loci in different continents reveal the existence of both ascertainment bias and population-specific variation (Figure S2). Asymmetries in joint site frequency spectrum plots arise because genetic associations can only be detected for markers that are polymorphic in the discovery population.^48^ In Figure S2, European ascertainment yields an excess of SNPs on the left and right sides of panels A and B and African ascertainment yields an excess of SNPs on the top and bottom sides of panel C. Notably, 34% of the variants in the Hagenaars et al. PGS and 21% of the variants in the Heilmann-Heimbach et al. PGS are near monomorphic in Africa (MAF < 0.01). Similarly, 54% of independent African associations implicated in this present study are near monomorphic in Europe (MAF < 0.01).

While previous studies have noted that Neanderthal introgression impacts genes that affect skin and hair morphology-related traits in European genomes,^24,49,50^ none so far have tested for a direct association between introgressed alleles and MPB. Importantly, African genomes are largely free of Neanderthal DNA. This difference between European and African genomes could be a potential source of divergence in the genetic architecture of MPB. Hence, we tested for enrichment for Neanderthal introgression in autosomal baldness-associated variants that were ascertained in European and African populations. Using Skov’s introgression map,^43^ we calculated the introgression frequencies of Neanderthal DNA in non-overlapping 10 kb windows around baldness-associated SNPs and compared these frequencies against 1,000 sets of matched control SNPs. If Neanderthal introgression is a driver of differences in the genetic architecture of MPB we would expect to find that European-ascertained baldness loci are enriched for Neanderthal DNA compared to the rest of the genome. However, neither the Hagenaars et al. PGS nor the Heilmann-Heimbach et al. PGS was enriched for Neanderthal DNA (Figure 4A-B). These results indicate that continental differences in the genetic architecture of MPB are not due to the differential introgression of Neanderthal alleles.

**Figure 4.**
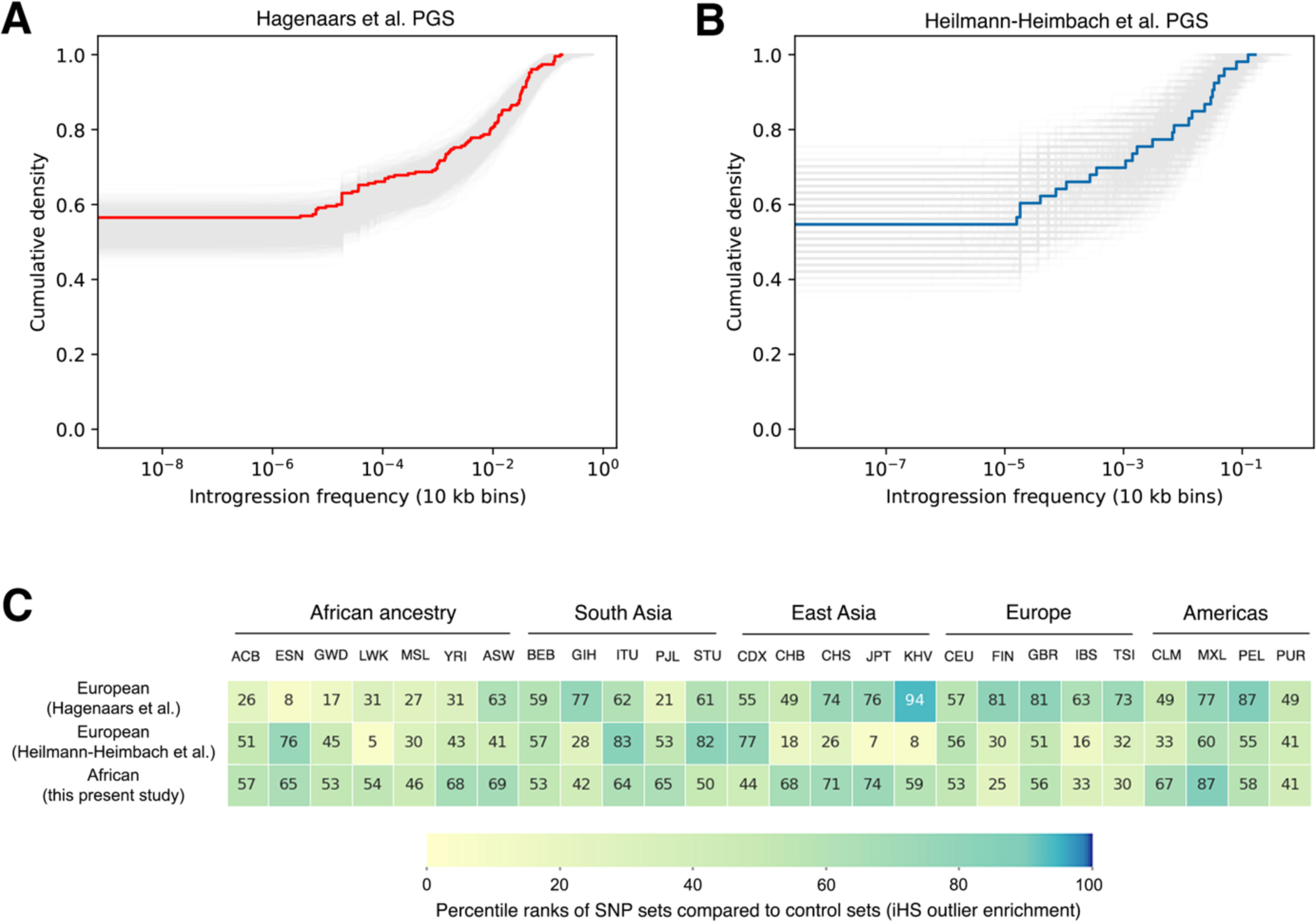
Tests of enrichment of Neanderthal DNA and recent polygenic selection. Cumulative densities of baldness-associated SNPs were compared to 1,000 sets of matched control SNPs, shown in gray. Most genomic loci have low introgression frequencies. (A) Loci from the Hagenaars et al. PGS, shown in red, are not enriched for introgressed Neanderthal DNA. (B) Loci from the Heilmann-Heimbach et al. PGS, shown in blue, are not enriched for introgressed Neanderthal DNA. (C). Tests of iHS outlier enrichment for baldness-associated SNP sets (1KGP data). Percentile ranks were generated from comparisons with 1,000 sets of matched control SNPs. Overall, autosomal SNPs that are associated with MPB are not enriched for signatures of MPB.

Natural selection can contribute to poor portability of polygenic predictions.^51^ If local adaptation has been a driver of differences in the genetic architecture of MPB then we would expect baldness-associated loci to be enriched for outliers of selection scans. Here, we tested for evidence of polygenic selection by examining whether sets of baldness-associated alleles have greater extended haplotype homozygosity than sets of control SNPs in 26 global populations from the 1KGP. Regardless of ascertainment scheme, baldness-associated SNPs were not enriched for outlier iHS statistics compared to the rest of the genome (Figure 4C). These results indicate that differences in the genetic architecture of MPB have been governed more by neutral evolution than by natural selection. However, we note that our polygenic tests of selection focused on autosomal loci and that individual loci may be exceptions to the general pattern.

To further understand the source of variation in the genetic architecture of MPB, we investigated whether individual baldness-associated loci have divergent allele frequencies between Europe and Africa. Highly divergent loci were identified by calculating F_ST_ statistics for baldness-associated SNPs that were ascertained in European and African populations. Consistent with the polygenic scans of selection described above, most baldness-associated SNPs have F_ST_ statistics that resemble the rest of the genome (Figure 5). Although multiple outliers exist, this need not indicate that baldness-associated SNPs are direct targets of natural selection, as genetic hitchhiking can cause allele frequencies to differ greatly across populations. High F_ST_ SNPs from European studies of baldness include rs4649041 at 1p36, rs13092705 at 3q26, rs17053607 at 4q32, rs9300169 at 12p12, and three SNPs at Xq12 (rs5965561, rs12558842, and rs2497911). High F_ST_ SNPs from our African GWAS include rs143451223 at 1q41 and rs1204041 at Xq12.

**Figure 5.**
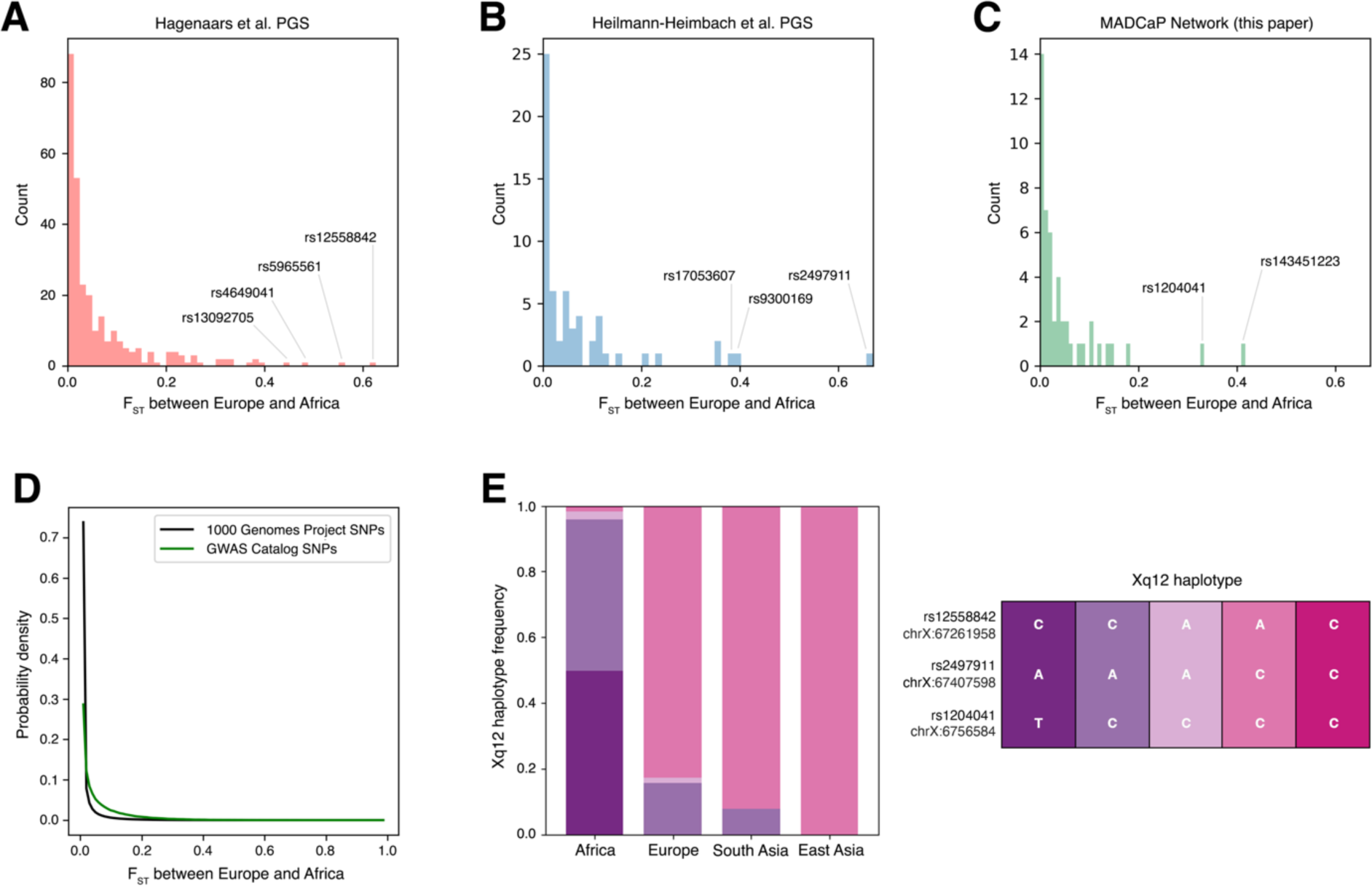
F_ST_ distributions and haplotype frequencies. Higher F_ST_ statistics are indicative of greater amounts of population structure, i.e., larger allele frequency differences between Europe and Africa (1KGP data). (A) Distribution of F_ST_ statistics of baldness-associated loci from the Hagenaars et al. PGS. (B) Distribution of F_ST_ statistics of baldness-associated loci from the Heilmann-Heimbach et al. PGS. (C) Distribution of F_ST_ statistics of African baldness-associated loci. (D) Genome-wide distributions of F_ST_ statistics from the 1KGP and SNPs in the NHGRI-EBI GWAS Catalog (all traits). (E). Haplotype frequencies at Xq12 differ across the globe. Genomic positions refer to build hg38, and the ancestral haplotype is AAC.

Multiple X-linked variants that are associated with MPB are evolutionary outliers.^52,53^ Key associations near the *EDA2R* and *AR* genes are rs12558842 in Hagenaars et al., rs2497911 in Heilmann-Heimbach et al., and rs1204041 in our study. Although the derived A allele at rs12558842 is nearly fixed in East Asian genomes, it is uncommon in Africa (1KGP allele frequencies of 0.998 in East Asia, 0.836 in Europe, and 0.060 in Africa). Similarly, the C allele at rs2497911 has an allele frequency of 0.998 in East Asia, 0.827 in Europe, and 0.041 in Africa. By contrast, the derived T allele at rs1204041 is common in Africa and all but absent from Eurasian populations (1KGP allele frequencies of 0.488 in Africa, 0.002 in Europe, and 0.000 in East Asia). The population-specificity of this allele is indicative of a recent evolutionary origin, i.e. the C ➝ T mutation at rs1204041 likely occurred after the out-of-Africa migration. These three variants are within 304kb of each other and in linkage disequilibrium, and additional insights can be gleaned from haplotype frequencies at Xq12 in different continental populations. We note that that the ancestral AAC Xq12 haplotype is all but absent from global populations (Figure 5E). Intriguingly, multiple divergent haplotypes have risen to high frequency, including CAT in Africa and ACC in Eurasia (Figure 5E). These patterns are consistent with the Xq12 region having undergone multiple independent selection events. Intriguingly, the *EDA2R* gene at Xq12 is functionally related to the *EDAR* gene at 2q13, which has previously been associated with scalp hair thickness and implicated in a genome-wide scan of selection.^54^ The *AR* gene at Xq12 encodes a steroid-hormone activated transcription factor that has many downstream effects.^55^ We note that Xq12 is located near the centromere in a genomic region with low recombination rates. Because of genetic hitchhiking, divergent haplotype frequencies at Xq12 need not have been caused by selection acting directly on baldness.

## Discussion

Our analyses of a novel African dataset revealed that genetic predictions of MPB generalize poorly across continental populations. Although differences in the genetic architecture of this complex trait are likely the primary cause of the poor portability of polygenic predictions, one additional contributing factor is that the UK Biobank cohort has a median age of 57, while phenotypes in the MADCaP Network dataset refer to age 45. In the first GWAS of MPB in sub-Saharan Africa, we identified 51 marginally significant baldness associations, many of which are due to polymorphisms that are only found in Africa. Examining the evolutionary genetics of MPB, we found that private alleles contribute more to the poor portability of genetic predictions than population-specific introgression of Neanderthal DNA. In general, we found that continental differences in the genetic architecture of MPB are more due to neutral evolution than recent positive selection. However, exceptions to this pattern involve X-linked loci.

The complex evolutionary history of X-linked variants near the *EDA2R and AR* genes has implications for the prediction of androgenetic alopecia in different populations. We note that rs12558842 has the largest effect size in the Hagenaars et al. PGS and rs2497911 has the largest effect size in the in Heilmann-Heimbach et al. PGS,^26,28^ while the African MPB association that was implicated in our study (rs1204041) has only a modest effect size (Table S2). The contribution of a SNP to the genetic variance of a trait is maximized at intermediate allele frequencies, and the minor alleles at rs12558842 and rs2497911 are more common in Europe than in Africa. This implies that the impact of X-linked variants on MPB is greater in European populations than African populations. Knowing whether someone’s maternal grandfather was bald is more likely to be informative to European men than African men.

This study adds to the growing body of evidence that genetic predictions generalize poorly across populations. Although GWAS can yield valuable insights about the genetic architectures of complex traits,^11,56–58^ differences in genetic architecture have implications for downstream applications. Because approaches like Mendelian randomization are only effective at inferring causal relationships if genetic predictors work well, it is necessary to use well-tuned genetic instruments.^59,60^ Similarly, although some authors have suggested that PGS for traits like baldness may be useful in crime scene investigations and reconstructing the phenotypes of archaic humans,^61^ care must be taken when doing so. Our findings underscore the limitations of assuming that the genetic architectures of complex traits are the same across individuals and populations.

Our study of androgenetic alopecia is not without its limitations. One caveat is that sample sizes were relatively limited, which means that some of the marginally significant associations identified in this study are likely to be false positives. Nevertheless, this marks the first African study of its kind, and it provides important insight into how the genetic architectures of a complex trait can differ across populations. The pattern and degree of baldness was obtained by self-reported questionnaire using the Norwood-Hamilton baldness scale. Although this instrument has been widely used in various European population there is a possibility that it results in misclassification for African men. In addition, men included in this study were controls recruited from different hospital departments and there is potential that some of the comorbidities that they were hospitalized for may have had a potential impact on the baldness score and pattern. An additional complication is that the relatively poor portability of polygenic predictions may be due in part to genotype-environment interactions. We note that the study sites analyzed here contain individuals from a wide range of sociolinguistic groups and genetic ancestries, and there may substantial heterogeneity in the genetic architecture of MPB and other complex traits within Africa.^62,63^ Going forward, there is clear need to conduct additional studies of diverse African populations (including functional genomics experiments to infer molecular mechanisms that are due to private alleles). Because early-onset MPB is indicative of the levels of sex hormones,^64^ future studies combining baldness phenotypes with genetic data may be informative about the risks of prostate cancer and other hormonal diseases.

## Data and code availability

GWAS summary statistics are available by request at https://madcapnetwork.org/. Genotype data presented in this analysis are publicly available via dbGAP (accession: phs002718.v1.p1). Additional data are available through controlled access upon request through the MADCaP network.

## Supplemental information

Supplemental information includes three supplemental tables and two supplemental figures.

## Supporting information

Table S2. Summary statistics for the 51 African associations with MPB.

Table S3. Pathway enrichment statistics.

## Acknowledgements

We thank the individuals who participated in this study. This work was supported by NIGMS Grant R35GM133727. This study is a product of the MADCaP Network (NCI grants U01CA184374 and R01CA257328).

## Author contributions

Supervised research: JL. Conceived and designed the experiments: IA and JL. Performed experiments: PK and LNP. Performed statistical analysis and analyzed data: RJ, UH, AP, MH, MSK, ME, and WCC. Contributed reagents/materials/analysis tools: MoJ, JEM, AAA, BA, MaJ, SMG, OIA-S, PWF, and AOA. Wrote the paper: RJ, UH, AP, AO, TER, IA, TRR, AOA, IA, and JL. Other (funding acquisition): JL and TRR and JL. Other (project administration): CA.

## Web resources

1000 Genomes Project (1KGP): https://www.internationalgenome.org/

Hagenaars et al. PGS: https://doi.org/10.1371/journal.pgen.1006594.s002

Heilmann-Heimbach et al. PGS: https://www.nature.com/articles/ncomms14694

LDlink: https://ldlink.nih.gov/

MADCaP Network: https://madcapnetwork.org/

NHGRI-EBI GWAS Catalog: https://www.ebi.ac.uk/gwas/

POLMM: https://github.com/WenjianBI/POLMM/

Skov’s introgression map: http://tinyurl.com/5dbwfpvk

snpXplorer: https://snpxplorer.net/

## Supplemental tables

**Table S1.**
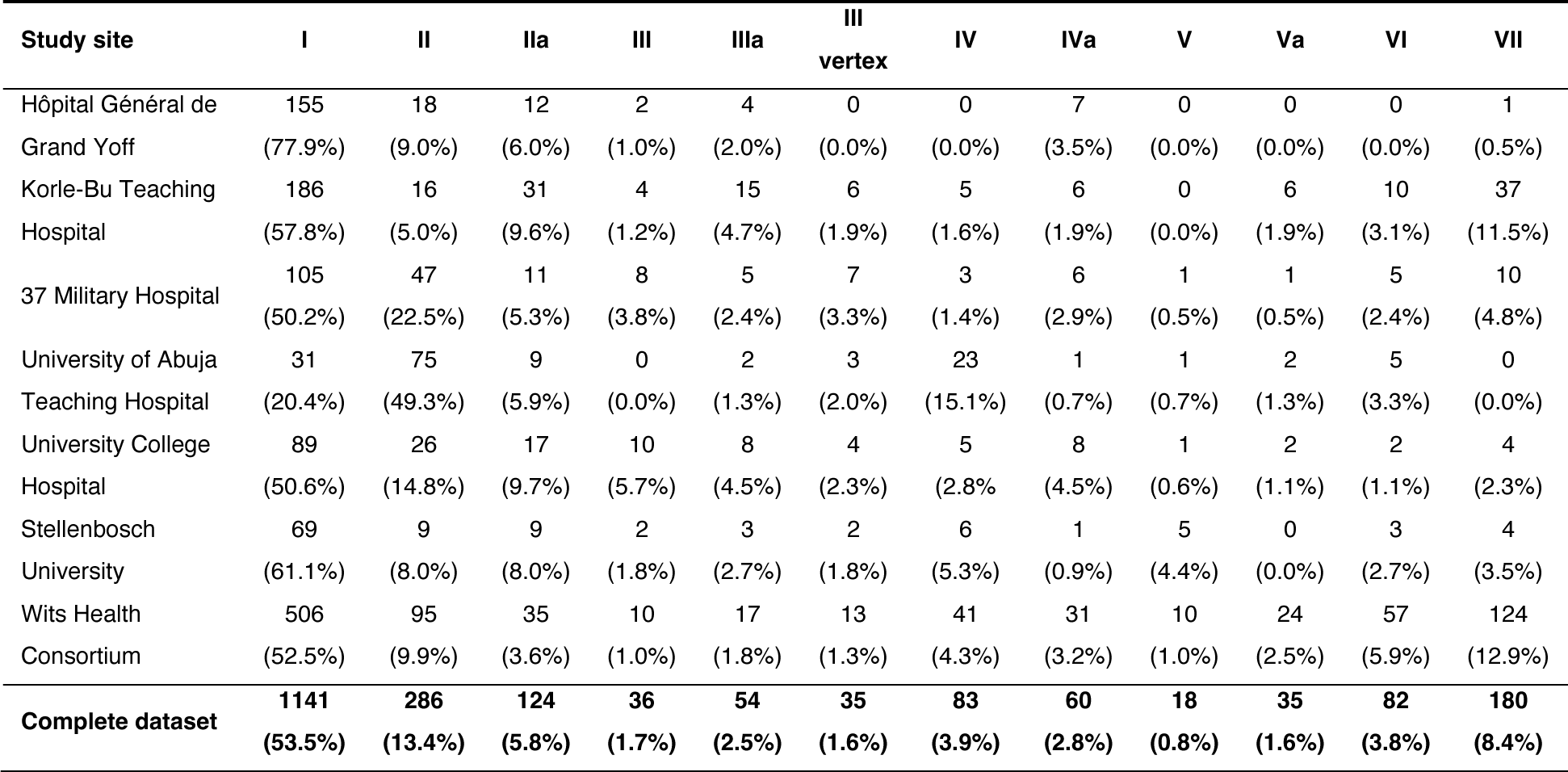
Hamilton-Norwood baldness patterns at each African study site. Self-reported baldness scores at age 45 are reported here.

**Table S2. Summary statistics for the 51 African associations with MPB.** This table lists SNP IDs, genomic positions, GWAS summary statistics, whether African baldness hits are in LD with previously published baldness hits, and GTEx annotations. Population genetics statistics include allele frequencies of African controls (i.e., non-bald individuals) and 1KGP populations. F_ST_ statistics (including genome-wide F_ST_ percentiles), and iHS outlier percentile ranks in 26 populations from the 1KGP. This supplemental table is available as a separate tab-delimited file.

**Table S3. Pathway enrichment statistics.** Gene-set enrichment analyses were performed using snpXplorer.^40^ P-values are false discovery rate-corrected for GO:BP (FDR: 1%) and Reactome (FDR: 10%) pathways. This supplemental table is available as a separate tab-delimited file.

**Figure S1.**
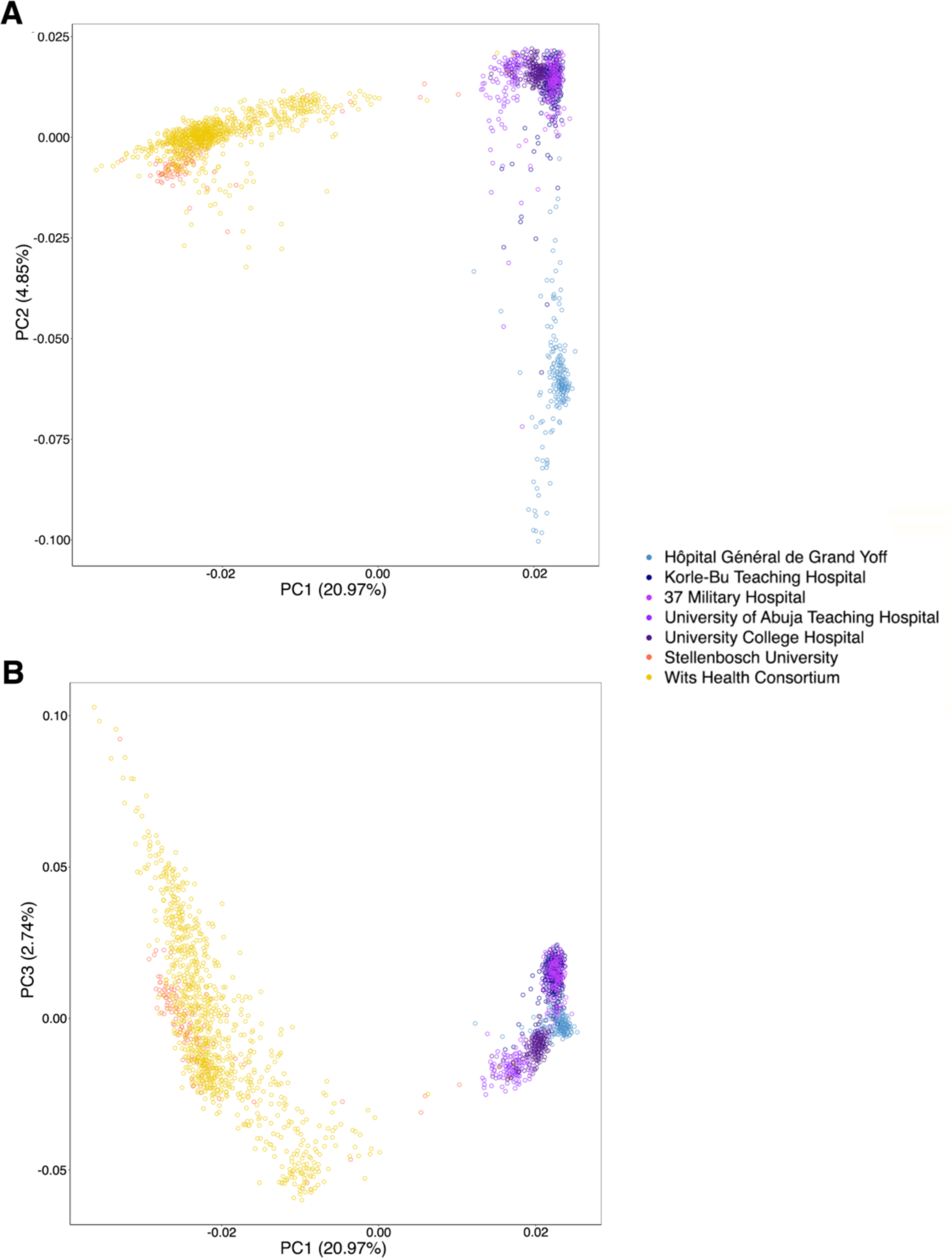
African population structure. PCA of the 2,136 MADCaP Network samples analyzed here. Data points are color coded by study site: Senegalese samples are represented by blue points, purple Ghanaian and Nigerian samples are represented by purple points, and South African samples are represented by orange points. (A) PC1 vs. PC2. (B) PC1 vs. PC3.

**Figure S2.**
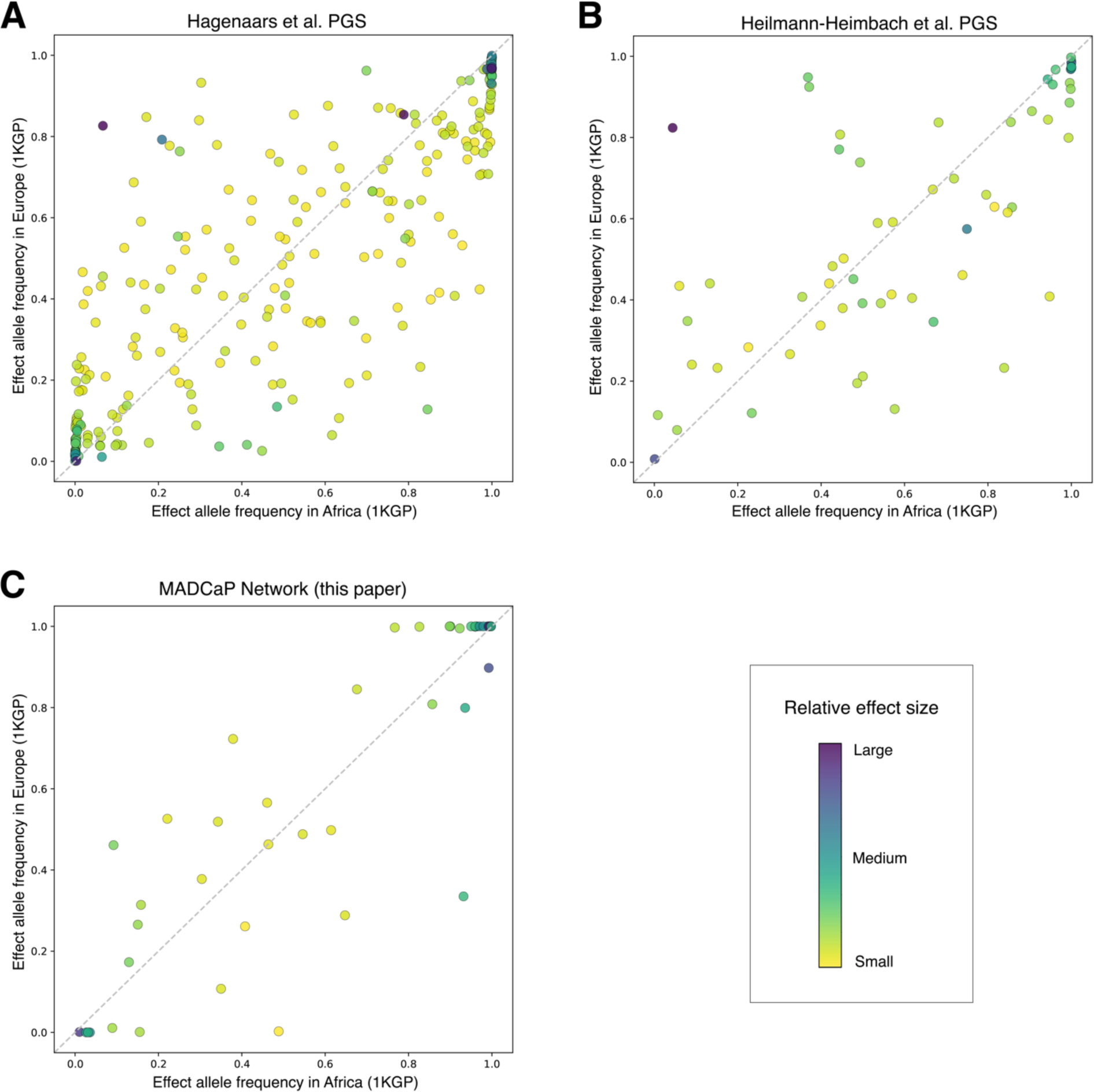
Allele frequencies of genetic variants that are associated with androgenetic alopecia. Joint site spectra show 1KGP data from Europe and Africa. Coloration indicates the relative effect size of each MPB-associated variant. (A) Allele frequencies of European-ascertained baldness associations from the Hagenaars et al. PGS. (B) Allele frequencies of European-ascertained associations from the Heilmann-Heimbach et al. PGS. (C) Allele frequencies of African-ascertained associations from this present study.

